# Compartment-Specific Antibody Correlates of Protection to SARS-CoV-2 Omicron in Macaques

**DOI:** 10.1101/2024.03.01.582951

**Authors:** Xin Tong, Qixin Wang, Wonyeong Jung, Taras M. Chicz, Ross Blanc, Lily J. Parker, Dan H. Barouch, Ryan P. McNamara

## Abstract

Antibodies represent a primary mediator of protection against respiratory viruses such as SARS-CoV-2. Serum neutralizing antibodies (NAbs) are often considered a primary correlate of protection. However, detailed antibody profiles including characterization of antibody functions in different anatomic compartments are not well understood. Here we show that antibody correlates of protection against SARS-CoV-2 challenge are different in systemic versus mucosal compartments in rhesus macaques. In serum, neutralizing antibodies were the strongest correlate of protection and were linked to Spike-specific binding antibodies and other extra-neutralizing antibody functions that create a larger protective network. In contrast, in bronchiolar lavage (BAL), antibody-dependent cellular phagocytosis (ADCP) proved the strongest correlate of protection rather than NAbs. Within BAL, ADCP was linked to mucosal Spike-specific IgG, IgA/secretory IgA, and Fcγ-receptor binding antibodies. Our results support a model in which antibodies with different functions mediate protection at different anatomic sites. The correlation of ADCP and other Fc functional antibody responses with protection in BAL suggests that these antibody responses may be critical for protection against SARS-CoV-2 Omicron challenge in mucosa.

## Introduction

COVID-19 vaccines, which generate antibodies to the severe acute respiratory syndrome coronavirus 2 (SARS-CoV-2) Spike protein, have shown remarkable success at attenuating severe disease. Neutralizing antibodies, which most commonly target the receptor binding domain (RBD) of Spike, were identified as a correlate of protection against ancestral strains of SARS-CoV-2 ^1^. However, as Omicron-lineage SARS-CoV-2 variants emerged, vaccine-/infection-acquired antibody neutralization was largely lost due to the high degree of antigenic shift within the RBD ^2-6^. Yet protection from disease in vaccinated individuals did not see a concomitant drop ^7-11^, signifying that immune mediators of protection other than neutralizing antibodies existed.

Beyond their capacity to neutralize, antibodies exert several non-neutralizing functions such as antibody-dependent opsinophagocytosis, antibody-dependent cellular cytotoxicity, and complement deposition ^12^. These functions are largely modulated by post-translational modifications to the crystallizable fragment (Fc) of antibodies which dictate their binding to Fc-receptors (FcγR for IgG subclasses, FcαR for IgA subclasses, etc.) on the surface of immune cells. Previous reports have demonstrated that FcγR-binding antibodies can recognize highly diverged SARS-CoV-2 Spikes and confer protection even when neutralization is lost ^13-15^. To that end, antibodies mediate protection against pathogens such as SARS-CoV-2 through a variety of functions.

It is unclear how antibody correlates of protection for COVID-19 are shaped in different anatomic compartments. In this study, we show the unexpected results that neutralizing antibodies are a strong correlate of protection in serum, but are not a clear correlate of protection in mucosa. Instead, extra-neutralizing functions such as antibody-dependent cellular phagocytosis were the strongest correlate of protection in bronchiolar lavage (BAL) against SARS-CoV-2 Omicron challenge. This extra-neutralizing role was conserved across SARS-CoV-2 variant Spikes, including the challenge strain. Our results support a model in which antibody correlates of protection against SARS-CoV-2 Omicron are different in different anatomic compartments.

## Methods

### Experimental Animals and Study Design

This study in rhesus macaques has been published previously and serum and BAL were collected for secondary use ^16^. Briefly, 4-8 year old *Rhesus macaques* were administered two doses of Ad26.COV2.S vaccine IM and one IM boost with either Ad26.COV2.S or Ad26.COV2.351 (Beta Spike). The NHPs were given boosters with a bivalent Ad26.COV2.S + Ad26.COV2.S.529 (BA.1 Spike) by IM, IN, or mucosal (intratracheal) route (6-8 animals/group). Serum and BAL collections were done at week 0 (pre-boost), week 4, and week 15 post-boost. Animals were subsequently challenged at week 16 with SARS-CoV-2 Omicron BQ.1.1 with 2E+6 PFU through intratracheal delivery. Viral loads were monitored and quantified in the lower respiratory tract (Supplementary Figure 1A). As described in the previous study ^16^, all animal study protocols were designed and conducted in compliance with all relevant local, state, and federal regulations and were approved by the Bioqual Institutional Animal Care and Use Committee (IACUC).

### Ig Subclassing/Isotyping and FcγR binding

Levels of antigen-specific antibody subclasses/isotypes and Fc-gamma receptor (FcγR) interaction were evaluated via the multiplexing Luminex microsphere-based assay, as previously described ^17^. Antigens of target were covalently linked to carboxyl group-labeled MagPlex microspheres (Luminex) through NHS-ester linkages using Sulfo-NHS and EDC (Thermo Fisher). Serum and BAL samples were diluted (serum isotypes/subclasses and FcγR binding: 1:250, BAL isotypes/subclasses and FcγR binding: 1:25) and added to the antigen-coupled microspheres to form the immune complexes in 384-well plates, and subsequently incubated at 4°C overnight, shaking at 750 rpm. After incubation, plates were washed with the washing buffer containing 0.1% BSA and 0.02% Tween 20 in PBS. Following the wash step, antibody isotype/subclass-specific mouse anti-rhesus antibodies (NHP Reagent Resource) were added to the immune complexes and incubated at room temperature for 1 h. Following a second wash step, the anti-mouse IgG Fc cross-adsorbed secondary antibody (PE, Thermo Fisher) was added to detect the anti-rhesus antibodies with fluorescence. For measurement of FcγR binding activities, Avi-tagged *Rhesus macaque* FcγRs (Duke Human Vaccine Institute) were biotinylated using BirA500 kit (Avidity) per manufacturer’s instructions and tagged with streptavidin-PE. The PE-labeled FcγR was subsequently incubated with the immune complexes for 2 h at room temperature. The plates were then washed and subject to Flow Cytometry measurements (iQue, Intellicyt) to determine the median fluorescence intensity (MFI). All Luminex experiments were conducted in duplicate, and the final results reported show the average values of the duplicates. The reagents and materials used are listed in the Key Resources Tables.

### Antibody-dependent Cellular Phagocytosis and Neutrophil Phagocytosis

ADCP and ADNP experiments were performed as previously described ^18,19^. Briefly, antigen proteins of the target were biotinylated using the EZ-link™Sulfo-NHS-LC-LC-Biotin kit (Thermo Fisher), then coupled to the fluorescent neutravidin beads (Thermo Fisher, F8776). The bead-antigen conjugates were incubated with diluted serum and BAL samples (serum: 1:100, BAL: 1:10) for 2 h at 37°C. The unbound antibody was removed by washing buffer. The immune complexes were then incubated overnight with cultured THP-1 cells (ADCP), or for 1 h with primary neutrophils isolated from human whole blood (ADNP) using negative selection (Stemcell). Treated THP-1 cells were subsequently washed and fixed in 4% paraformaldehyde (PFA), while the treated neutrophils were washed, stained for CD66b+ marker (Biolegend), and fixed in 4% (PFA) prior to flow cytometry analysis. A phagocytosis score for THP-1 or neutrophil was eventually determined as (% cells positive × Median Fluorescent Intensity of positive cells). Flow cytometry was performed with an iQue (IntelliCyt) instrument and population measurements were conducted using IntelliCyt ForeCyt (v8.1). The reagents and materials used are listed in the Key Resources Tables.

### Antibody-dependent Complement Deposition (ADCD)

ADCD assays were designed and performed as previously described ^20^. Antigens of target were covalently linked to the carboxyl group-labeled MagPlex microspheres (Luminex) through NHS-ester linkages using Sulfo-NHS and EDC (Thermo Fisher) as described for Luminex. Diluted serum and BAL samples (serum: 1:50, BAL: 1:10) were incubated with coupled antigens for 2 h at 37 °C to form immune complexes in 384-well plates. Plates were washed and incubated with lyophilized guinea pig complement (Cedarlane) diluted in gelatin veronal buffer with calcium and magnesium (Sigma Aldrich) for 20 min at 37 °C. The deposition of C3 complement component was evaluated by an anti-guinea pig C3 FITC detection antibody (MpBio). Fluorescent intensity was acquired using an iQue Flow Cytometer (Intellicyt). The antibody-specific complement C3 deposition is calculated as the median fluorescence intensity of FITC. All ADCD experiments were conducted in duplicate, and final values were reported as average of the duplicates. The reagents and materials used are listed in the Key Resources Tables.

### Antibody-dependent Natural Killer Cell (NK) Activation (ADNKA)

ADNKA assays were designed and performed as described previously ^21^. ELISA plates were coated 3 μg/mL of selected antigen and incubated at 4 °C overnight. The coated plates were washed with PBS and blocked with 5% bovine serum albumin (BSA) for 2 h at 37°C. Natural Killer (NK) cells were isolated from Leukopaks (Stemcell Technologies) using EasySep™ Human NK Cell Isolation Kit (Stemcell Technologies). The isolated NK cells were incubated overnight at 37°C 5% CO2 in R10 (RPMI-1640 (Sigma Aldrich) media supplemented with 10% fetal bovine serum (FBS) (Sigma Aldrich), 5% penicillin/streptomycin (Corning, 50 μg/mL), 5% L-glutamine (Corning, 4 mM), 5% HEPES buffer (pH 7.2) (Corning, 50 mM) supplemented with 2 ng/mL IL-15. The ELISA plates were washed, and diluted serum and BAL samples (serum: 1:40, BAL: 1:10) were added to plates for 2 h at 37°C to form immune complexes. After wash, NK cells were added to plates at a concentration of 2.5E+5 cells/mL in R10 media supplemented with anti-CD107a–phycoerythrin (PE)–Cy5 (BD Biosciences, lot # 0149826, 1:1000 dilution), brefeldin A (10 μg/mL) (Sigma-Aldrich), and GolgiStop (BD Biosciences). The NK cells were incubated with immune complexes for 5 h at 37°C. The incubated NK cells were stained for cell surface markers with anti-CD3 Pacific Blue (BD Biosciences, clone G10F5)), anti-CD16 allophycocyanin (APC)-Cy5 (BD Biosciences, clone 3G8), and anti-CD56 PE-Cy7 (BD Biosciences, clone B159) for 15 min at room temperature. The washed NK cells were then fixed with PermA (Life Technologies), permeabilized with PermB (Life Technologies), and labeled with anti-MIP-1β PE (BD Biosciences) and anti-IFNγ FITC for 15 min at room temperature. Fluorescent intensity was measured using iQue Cytometer (Intellicyt). NK cells were gated as CD56+/CD16+/CD3- and the NK activation was evaluated as the percentage of NK cells positive for CD107a, IFNγ, or MIP-1b. All assays were performed with at least two healthy donors and the results shown here report the average of the donors. The reagents and materials used are listed in the Key Resources Tables. A gating strategy figure can be found in Supplementary Figure 1B-H)

### Statistical analysis

All statistical analysis was performed with R (version 4.1.1). For Multivariate analysis, “systemsseRology” R package was used (https://github.com/LoosC/systemsseRology). The data were centered and scaled before the multivariate analysis. Partial Least Square Regression (PLSR) model was utilized to determine the best feature combination that describes peak viral load within the lower respiratory tract. Regression against peak viral loads was the primary study output, independent of route of booster delivery. Features that contribute most to the regression model were selected by LASSO (Least Absolute Shrinkage and Selection Operator). Features that were selected from more than 90% (Serum samples) or 80% (BAL samples) of 100 LASSO selections were finally selected for PLSR. PLSR model performance was determined by 5-fold cross-validations that were repeated 40 times. In addition, control models with random features or permuted output labels were built 25 times, whose accuracies were compared with the accuracy of the original model with 5-fold cross-validations. For LASSO-selected features were shown in the order of Variable Importance in Projection (VIP). In Figure 1D, the VIP score was calculated using a PLSR model that incorporates all measured features, to show the VIP score of neutralizing antibody titers that were not selected by LASSO. Features that are highly correlated with LASSO-selected features were plotted in network plots. Correlations with Spearman’s correlation > 0.6 with a false discovery rate (FDR) < 0.05 for serum and BAL samples.

**Figure 1.**
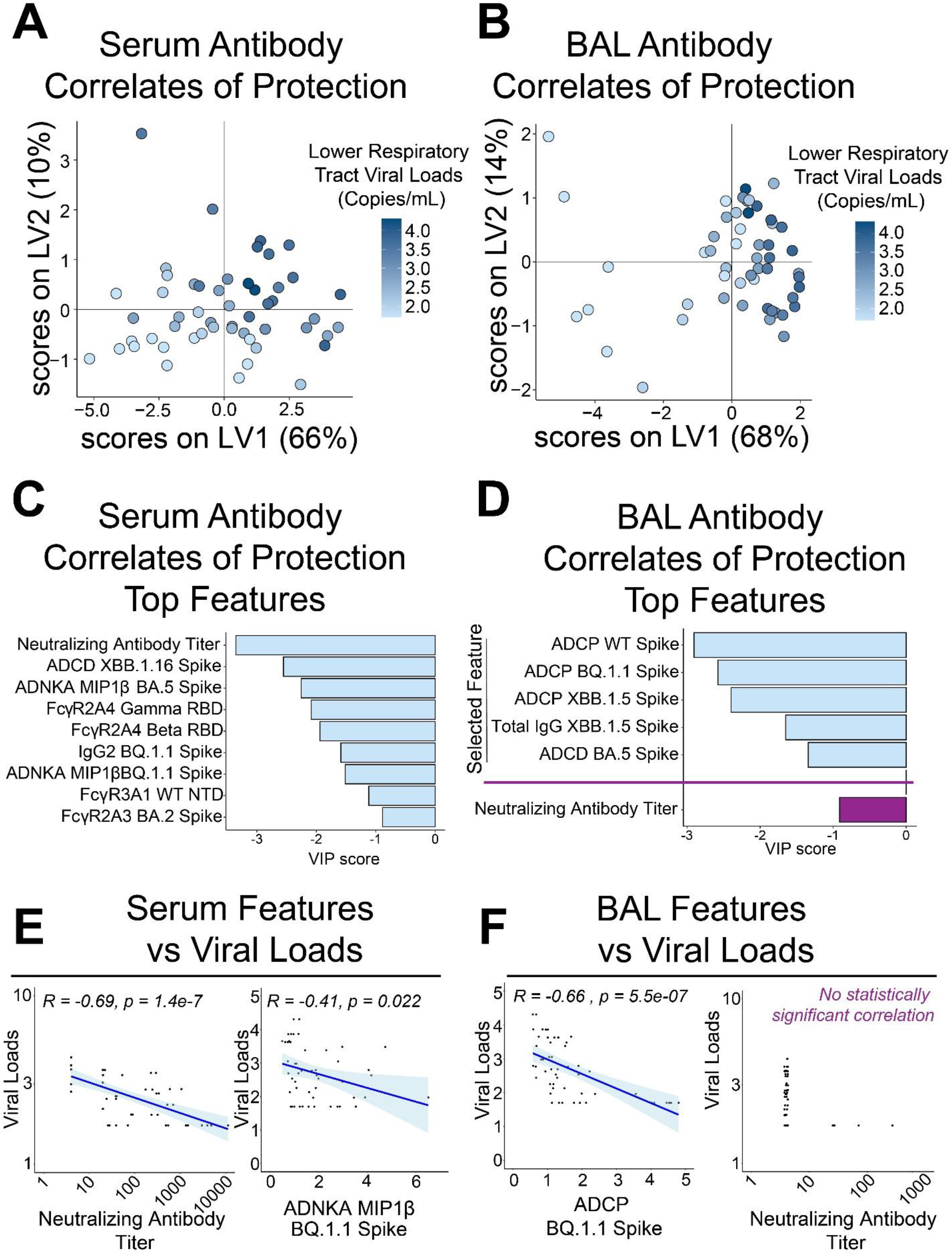
Defining antibody correlates of protection within serum and bronchiolar lavage (BAL). (A) Partial least squares regression (PLSR) model of serum antibody features of vaccinated and boosted NHPs inversely correlated with viral loads within the lower respiratory tract. Heatmap gradient of viral loads is shown on the right. (B) PLSR model of BAL antibody features of vaccinated and boosted NHPs inversely correlated with viral loads within the lower respiratory tract. Heatmap gradient of viral loads is shown on the right (C) Top serum antibody correlates of protection in the PLSR model. (D) Top BAL antibody correlates of protection in the PLSR model. Neutralizing antibody titer was not selected as a *bona fide* correlate in the BAL and was manually plotted in purple for comparison. (E) Validation of PLSR-selected serum correlates of protection. Viral loads were inversely correlated with (left) neutralizing antibody titer and (right) antibody-dependent natural killer cell activation (ADNKA) as measured by macrophage inflammatory protein 1 beta (MIP1β) production. Spearman’s R values and multiple comparisons adjusted p-values are shown. (E) Validation of PLSR-selected BAL correlates of protection. Viral loads were inversely correlated with (left) antibody-dependent cellular phagocytosis (ADCP) to BQ.1.1 Spike (challenge strain), but not to neutralizing antibody titers. Spearman’s R values and multiple comparisons adjusted p-values are shown only for the statistically significant ADCP.

For univariate analysis, the “rstatix” R package was used, and two-sided Wilcoxon tests were performed to determine if data from different timepoints (week 0 vs week 4, week 4 vs week 15, and week 0 vs week 15) significantly differed. P values were then corrected for multiple comparisons through FDR correction. Resulting p-values <0.05 were considered as significant. For all groups, baseline antibody levels were standardized to 1 to quantify fold changes. Moving averages from week 0, week 4, and week 15 post-booster were modeled and plotted showing the mean and 95% confidence intervals.

## Results

### Identification of humoral features inversely correlated with SARS-CoV-2 viral loads by anatomic compartment

Binding antibodies, Fcγ-receptor (FcγR) binding antibodies, neutralization titers, and Fc-effector functions from serum- and lower respiratory tract-resident antibodies were quantified through systems serology (Supplementary Figures 2-3). A composite multivariate probable least squares regression (PLSR) model was built to identify antibody features correlated with protection against viral loads across treatment groups. This was done for antibodies in serum (**Figure 1A**) and bronchiolar lavage (BAL) (**Figure 1B**). Viral loads scattered on the latent variable 1 axis (LV1), which accounted for 66% and 68% of the total variance explained in the serum and BAL humoral profiles, respectively.

Distinguishing antibody features driving protection by compartment were identified. In the serum, neutralizing antibody titers, FcγR-binding antibodies, antibody-dependent natural killer cell activations (ADNKA), and antibody-dependent cellular phagocytosis (ADCP) were all significant correlates of protection (**Figure 1C**, light blue bars). Interestingly, features driving protection in the BAL were focused on ADCP, IgG binding antibodies to Omicron XBB.1.5 Spike, and antibody-dependent complement deposition (ADCD) (**Figure 1D**, light blue bars). Since neutralizing antibodies were not selected as a driver of protection in BAL, we separately added it for comparison relative to the other features (**Figure 1D**, purple bar).

To confirm these results, we plotted correlations of selected features with viral loads based on compartments. As expected, serum selected features correlated with protection as defined by viral loads (**Figure 1E**, Supplementary Figure 4A). Likewise, BAL selected features correlated with protection (**Figure 1F**, left; Supplementary Figure 4B). No correlation between neutralizing antibody titer and viral loads in the BAL could be obtained (**Figure 1F**, right). PLSR models for both serum and BAL-resident selected features as driving protection were validated against permutated labels and random features (Supplementary Figure 5).

### BAL correlates of protection to SARS-CoV-2 are defined by binding, FcγR-binding, and effector functions

To investigate how the humoral landscape operates at the systems level, we created constellation linkage profiles by compartment. Features selected as bona fide correlates of protection by compartment were plotted with highly co-correlated features (see methods). Serum-resident antibody features driving protection were part of a large constellation of humoral features that included binding antibodies, FcγR-binding antibodies, and Fc-effector-mediated functions (Supplementary Figure 6).

The correlates of protection network of the BAL (site of infection) showed ADCP to BQ.1.1 Spike acting as a centralizing node linking to binding antibodies including IgG subclasses, IgA, secIgA, and FcγR-binding antibodies, as well as to other effector functions (**Figure 2A**). A separate correlate of protection in the BAL was total IgG, and a constellation of co-correlating features was mapped showing other binding IgG and FcγR-binding antibodies across VOC (**Figure 2B**). The last key correlate of protection identified was ADCD to XBB.1.5 Spike. As expected, this node correlated with ADCD to other Spike variants including Omicron sublineages (**Figure 2C**). The full constellation for BAL antibody correlates of protection is shown in Supplementary Figure 7.

**Figure 2.**
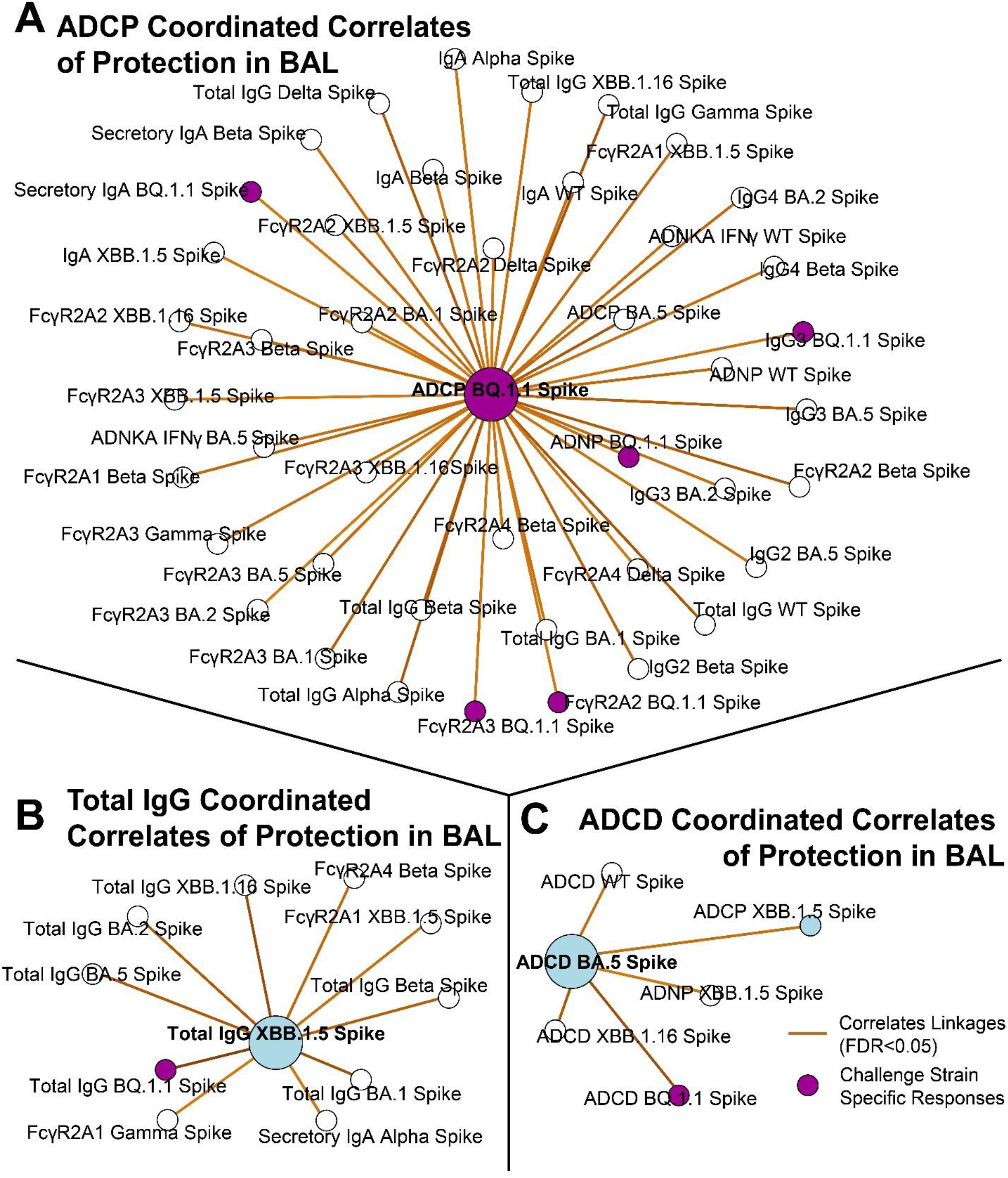
Antibody network of strongest correlates of protection in the BAL. A) ADCP to the challenge strain BQ. 1.1 Spike (purple node) was centered and correlating antibody features (R >0.6, FDR p-value <0.05) taken from the BAL are shown. B) Same as A, but for total IgG to XBB. 1.5 Spike. C) Same as A, but for ADCD to BA.5 Spike. Blue circles correspond to the selected features in Figure 1. Purple circles correspond to BQ.1.1 Spike, which was the challenge strain. Legend is shown at the bottom right.

Interestingly, while neutralizing antibodies were the strongest correlate of protection in the serum, neutralizing antibodies were neither a selected correlate of protection in the BAL nor were they linked to the identified correlates of protection.

### Mucosal boosting increases serum and lower respiratory tract resident humoral responses to divergent Spikes

To define if correlates of protection were influenced by route of vaccine booster, we analyzed the identified correlates of protection using the delivery site as a variable. For ADNKA responses, non-boosted NHPs showed no changes to any Spikes at any timepoints. IM-boosted (purple) NHPs showed significantly enhanced serum ADNKA to BQ.1.1 and XBB.1.5 Spikes. IN-boosted (orange) NHPs likewise showed enhanced ADNKA responses but were significant for BA.5 and XBB.1.5 Spike. Mucosal-boosted (blue) NHPs showed significant ADNKA responses to BA.5, BQ.1.1, and XBB.1.5 Spike (**Figure 3A**). Significant expansions of ADNKA to WT Spike were not observed for any groups, likely indicating that existing profiles to ancestral Spike from previous vaccinations were still present. Binding IgG within the serum to the Spikes strongly correlated with ADNKA. We thus assayed for total IgG binding to these Spikes from the serum of the boosted NHPs. We found that total IgG to Spike variants significantly increased for IM- and mucosal-boosted NHPs, but not for IN-boosted NHPs after multiple comparisons adjustments (**Figure 3B**).

**Figure 3.**
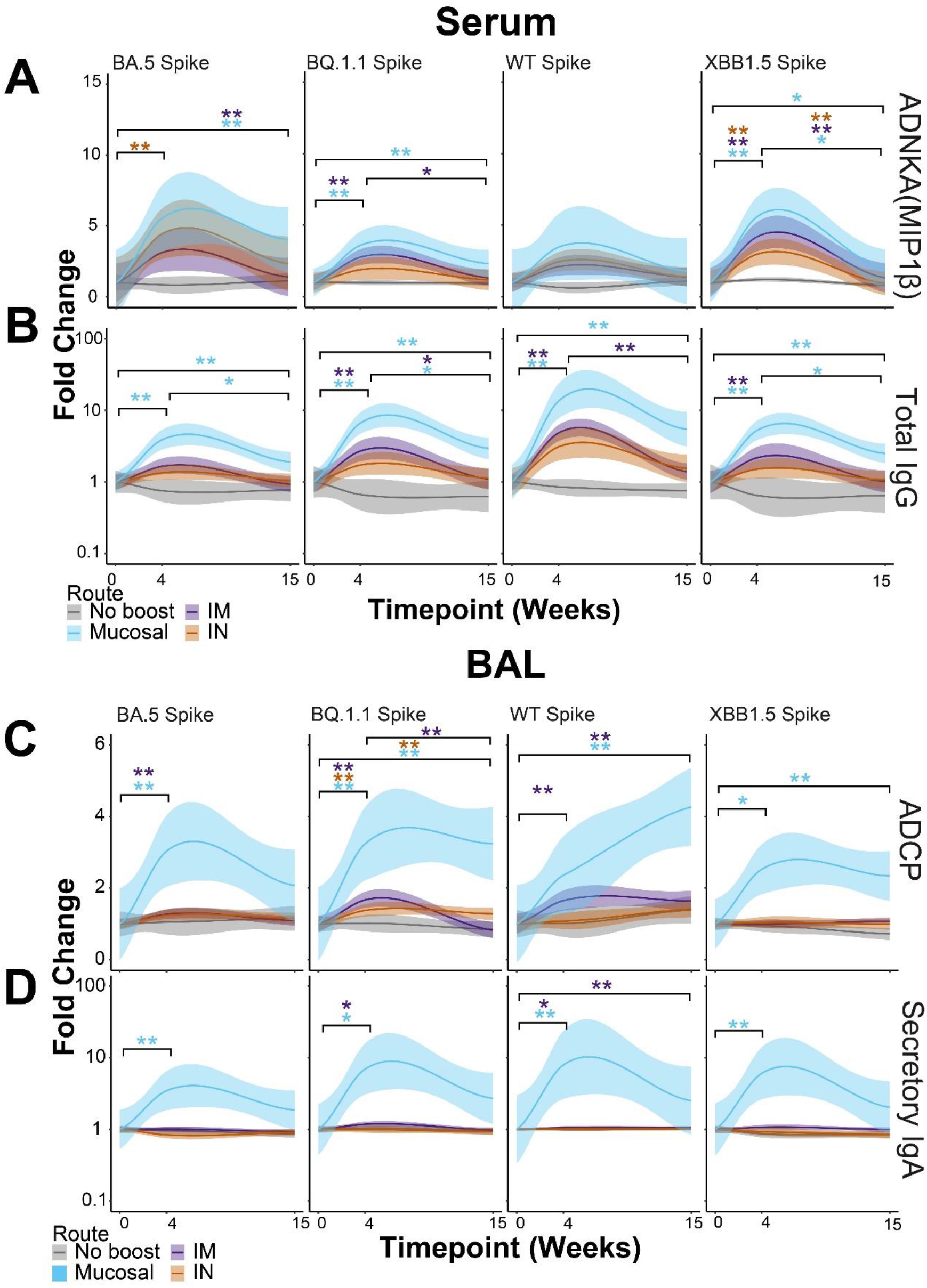
Mucosal boosting enhances serum and lower respiratory tract humoral responses to various SARS-CoV2 VoCs. A) Post-booster fold enhancements of serum-resident antibody-dependent natural killer cell activation (ANDKA) to the indicated Spike variants as quantified by macrophage inflammatory protein 1 beta (MIP1β) production. ADNKA was selected as a key correlate of protection in Figure 1. B) Same as A, but for the serum-resident networked feature of total IgG to the indicated Spikes variants. Color scheme legend is shown at the bottom for the serum responses. C) Post-booster fold enhancements of BAL-resident antibody-dependent cellular phagocytosis (ADCP) by monocytes to the indicated Spike variants. ADCP was selected as a key correlate of protection in Figure 1. D) Same as C, but for the BAL-resident networked feature of secretory IgA to the indicated Spike variants. Color scheme legend is shown at the bottom for the BAL responses. For all plots, fold enhancements were relative to the mean value of the group at the pre-boost time point, week 0. Shown here are the best-fit models of matched responses within 95% CI in regions shaded in corresponding colors. * = p < 0.05, ** = p < 0.01; Wilcoxon Test followed by FDR correction.

ADCP to several Spikes was the strongest correlate of protection in the BAL. Similar to serum responses, we analyzed how ADCP responses were shaped by booster delivery site. Interestingly, mucosal-boosted (blue) NHPs showed the strongest and most consistent ADCP to Spikes tested from BAL samples. IM-boosted (purple) NHPs also showed some significant ADCP expansions, but the magnitude of responses was much lower than mucosal-boosted NHPs. While some ADCP was observed from BAL samples from IN-boosted NHPs, only one Spike showed significance after multiple test corrections (**Figure 3C**). We last tested for secIgA responses in the boosted NHPs as it was strongly linked to ADCP within this compartment. Much to our surprise, secIgA was only found to be significantly induced in the BAL across Omicron Spikes for mucosal-boosted NHPs. This occurred at 4 weeks post-boost. IM-boosted NHPs showed low secIgA induction to some Spikes, while IN-boosted NHPs did not show any significant secIgA increases for any Spike assayed. Therefore the correlation between secIgA and the identified BAL correlates of protection was strongly driven by booster site.

Other serum and BAL humoral profiles showed similar results, including functional assays (Supplementary Figure 8-9). Within the BAL, mucosal-delivered boosts consistently yielded higher antibody levels and functional outputs. Collectively, these results confirm our machine learning approaches of classifying correlates of protection by compartment. These results are strongly influenced by the route of boosting.

## Discussion

In this study, we report that antibodies with distinct functions in different anatomic compartments are correlates of protection against SARS-CoV-2 Omicron challenge in rhesus macaques. Our data supports a model whereby mucosal antibody correlates of protection in the lower respiratory tract are primarily functional antibodies that mediate opsinophagocytosis, while serum antibody correlates of protection are primarily neutralization, binding, and non-neutralizing functions. Our data suggest that antibodies with Fc effector functions may be more important than currently appreciated at mucosal surfaces.

Previous work demonstrated that neutralizing antibodies were a correlate of protection against ancestral SARS-CoV-2 across several clinical trials ^1^. As SARS-CoV-2 adapted itself to the human population, VOCs emerged and neutralizing antibody capacity was progressively lost ^22-24^. This did not translate to a loss of clinical protection ^9-11^, indicating that correlates beyond neutralization existed. Previous reports have shown that FcγR-binding antibodies and their corresponding effector functions were required for protection against antigenically diverged Spikes ^13,25^.

We employed a systems serology approach to arrive at the conclusion that humoral profiles that correlate with protection are different in different anatomic compartments. Deep antibody profiling including a comprehensive analysis of antibody functions revealed that neutralizing antibody titers are the strongest correlate of protection in serum. This is in agreement with several reports showing that serum-resident neutralizing antibodies are strongly induced after vaccination and/or infection, and are a correlate of protection ^16,26-33^. Our approach extends these findings to show that antibody correlates of protection are different at distinct anatomic sites. We show that antibody correlates of protection at the mucosa are enriched for effector functions such as opsinophagocytosis. These extra-neutralizing functions were most tightly linked to mucosa IgG and IgA/secIgA. Stimulation of IgA by different COVID-19 vaccine formulations has been shown ^34^. Our study shows that IgA can be stimulated by distinct delivery sites of the same vaccine formulation. IgA is known to be a potent neutralizer ^35-37^ and a strong driver of opsinophagocytic function ^38^. Due to its polyfunctionality and mucosal localization, vaccine formulations and platforms have sought to enhance IgA responses ^34,39-42^. Our study supports the notion that vaccine delivery route can greatly impact mucosal IgA induction.

In conclusion, our study demonstrates that antibody correlates of protection may be different in different anatomic compartments. In the lower respiratory tract, antibody Fc effector functions leveraged by IgG and IgA drive protection against SARS-CoV-2, whereas in the serum, neutralizing antibodies drive protection. Further work characterizing how antibodies’ roles are influenced by their compartment can lead to vaccination strategies conferring multiple layers of protection. This is particularly important for emerging infectious diseases such as SARS-related coronaviruses.

## Supporting information

Key Resources Table

## Acknowledgments

We thank Terry and Susan Ragon and Mark and Lisa Schwartz for their contributions to the Ragon Institute of Mass General, MIT, and Harvard. This study was supported by the CA260476 (D.H.B.), the 1P01AI165072-01 (R.P.M.), the 2U19AI135995-06 (R.P.M.), and the Bill and Melinda Gates Foundation Global Health and Vaccine Accelerator Program (GH-VAP) INV-001650.

## Author Contributions

X.T. and Q.W. contributed equally as co-first authors to this manuscript. X.T., Q.W., D.H.B., and R.P.M. conceptualized the study. X.T., Q.W., T.M.C., R.B., and L.J.P. performed the wet-bench systems serology experiments. Q.W., W.J., and R.P.M. performed bioinformatic systems serology analyses. X.T., Q.W., W.J., D.H.B., and R.P.M. analyzed all the data and wrote the manuscript. D.H.B. and R.P.M. provided supervision and funding acquisition. All authors reviewed and edited the manuscript.

## Data Availability Statement

Raw data for the systems serology have been deposited on the RagonSystemSerology GitHub Homepage (https://github.com/RagonSystemSerology/SystSeroNHPCOP_ProjID20240222). Correspondence and requests for materials and resources should be addressed to R.P.M.

