## Supplementary Information for "Compartment-Specific Antibody Correlates of Protection to SARS-CoV-2 Omicron in Macaques"

### Supplemental Information

Supplemental Information contains Supplementary Figures 1-9 and corresponding Supplementary Figure Captions.

### Supplementary Figure 1

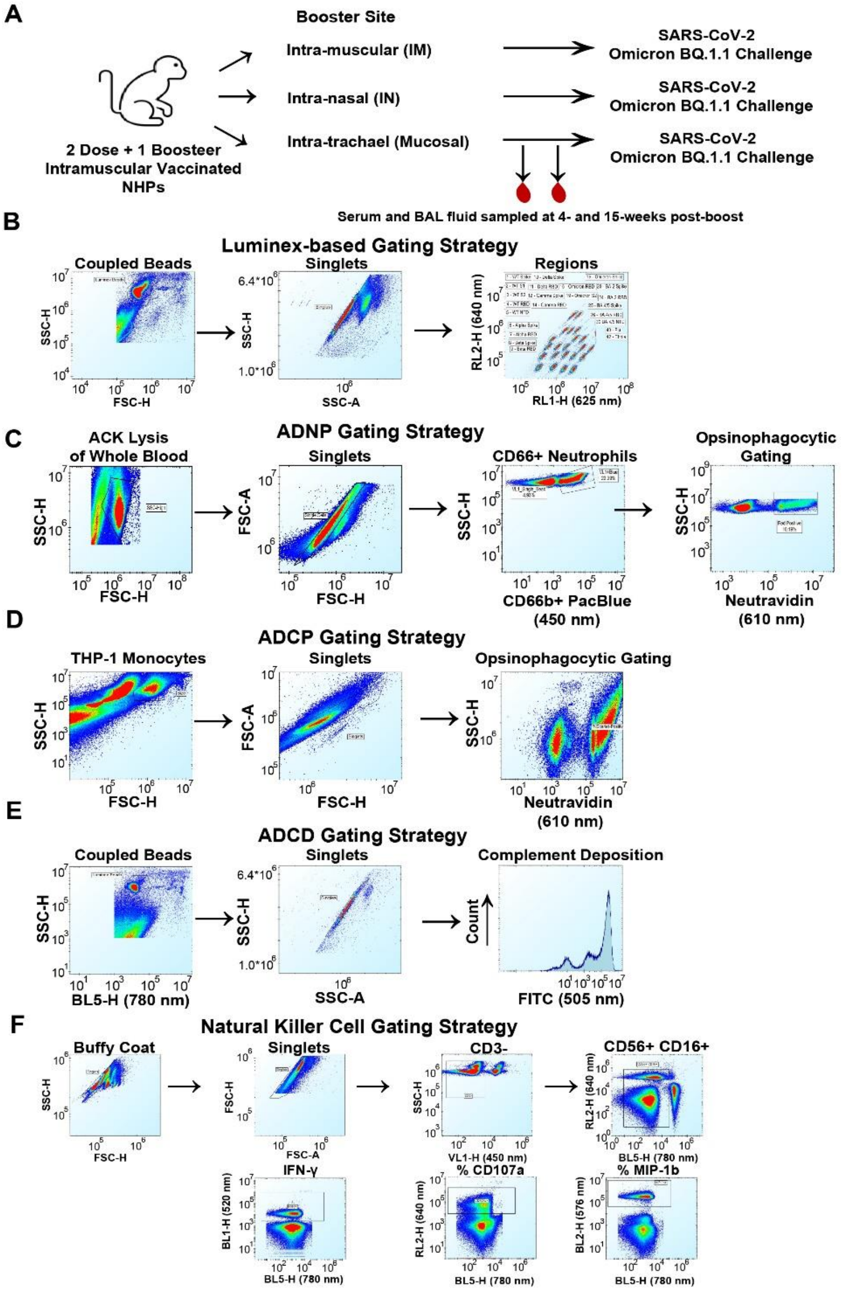


**Supplementary Figure 1. Cohort description and flow cytometry gating strategy.** A) Non-human primates primed with two-doses of intramuscular Ad26.COV2.S at week -108 and were boosted at week -69 with Ad26.COV2.S. At week 0, NHPs were boosted with a bivalent Ad26.COV2.S (ancestral Spike) + Ad26.COV2.S529 (Omicron BA.1 Spike) administered intra-muscularly (IM), intra-nasally (IN), or intra-trachael (mucosal). At weeks 4 and 15, serum and BAL samples were collected followed by challenge with SARS-CoV-2 Omicron BQ.1.1. B-F) Gating strategy for antibody-based B) binding profiles, C) antibody-dependent neutrophil phagocytosis (ADNP), D) antibody-dependent cellular phagocytosis by monocytes (ADCP), E)

antibody-dependent complement deposition (ADCD), and F) antibody-dependent natural killer cell activation (ADNKA).

### Supplementary Figure 2

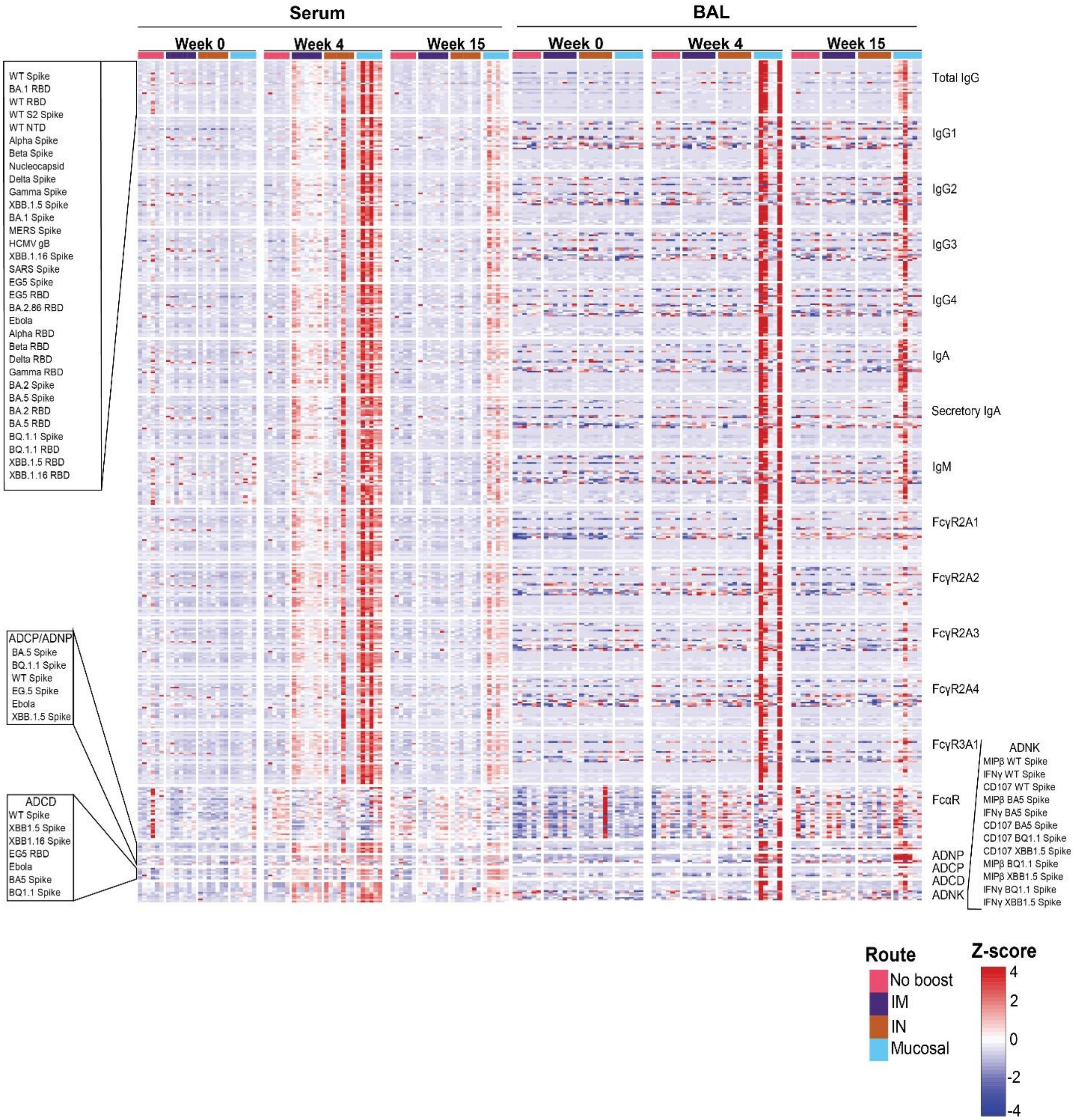


**Supplementary Figure 2. Full systems serology data array**. Serum (left) and BAL (right) antibody samples were assayed at week 0, week 4, and week 15 post-bivalent booster for binding to the antigens listed on the top left. Antibody subclass, isotype, Fc-binding, and functional outputs are shown on the right column. Route of administration and value legend is shown on the bottom right. All results were z-scored to account for differences in outputs.

**Supplementary Figure 3**

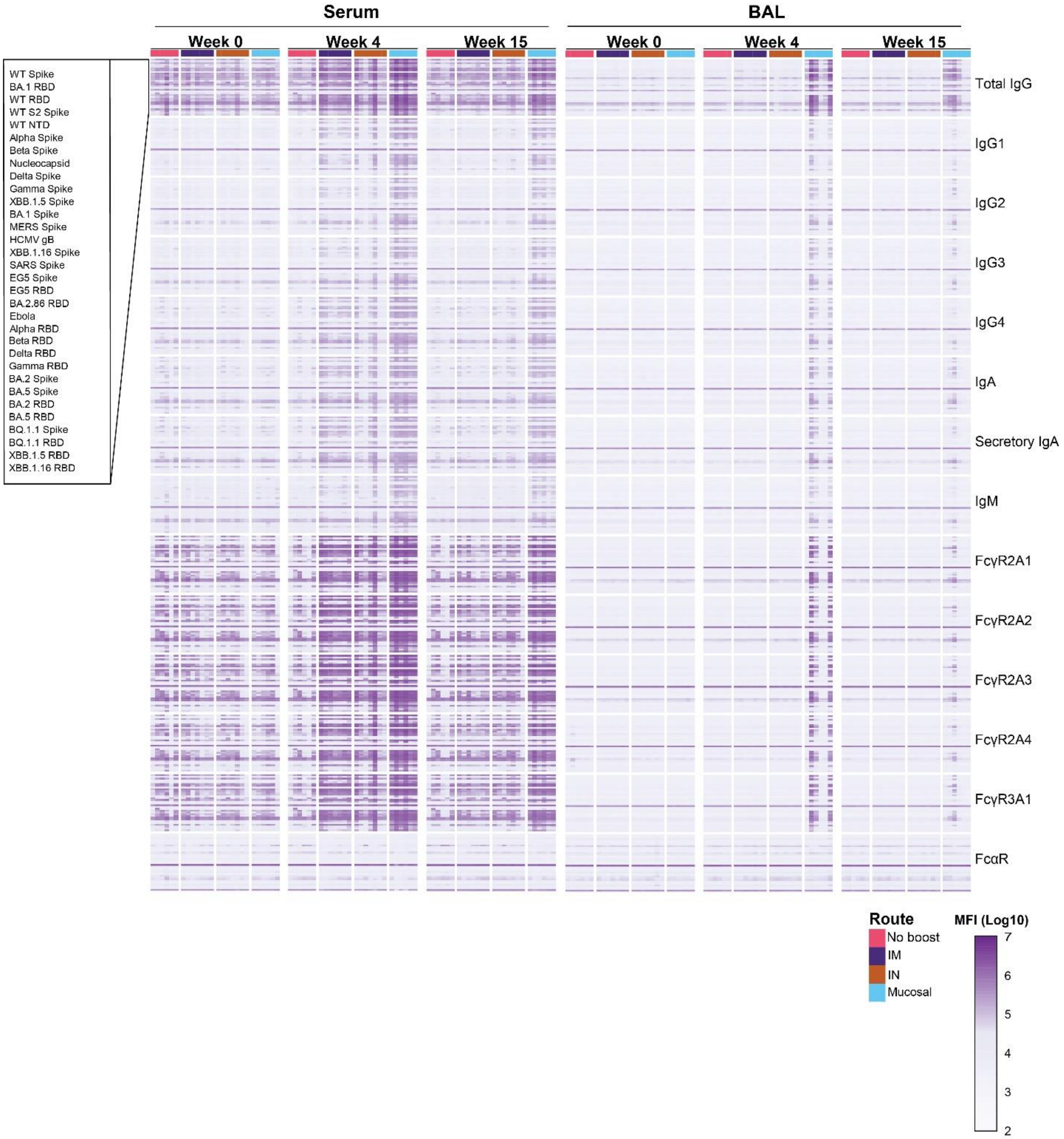


**Supplementary Figure 3. Biophysical antibody binding levels to various antigens in raw values**. Serum (left) and BAL (right) antibody samples were assayed at week 0, week 4, and week 15 post-bivalent booster for binding to the antigens listed on the top left. Antibody subclass, isotype, and Fc-binding antibodies are shown on the right column. Route of administration and value legend is shown on the bottom right. Binding levels are shown as a heatmap in log 10 values.

**Supplementary Figure 4**

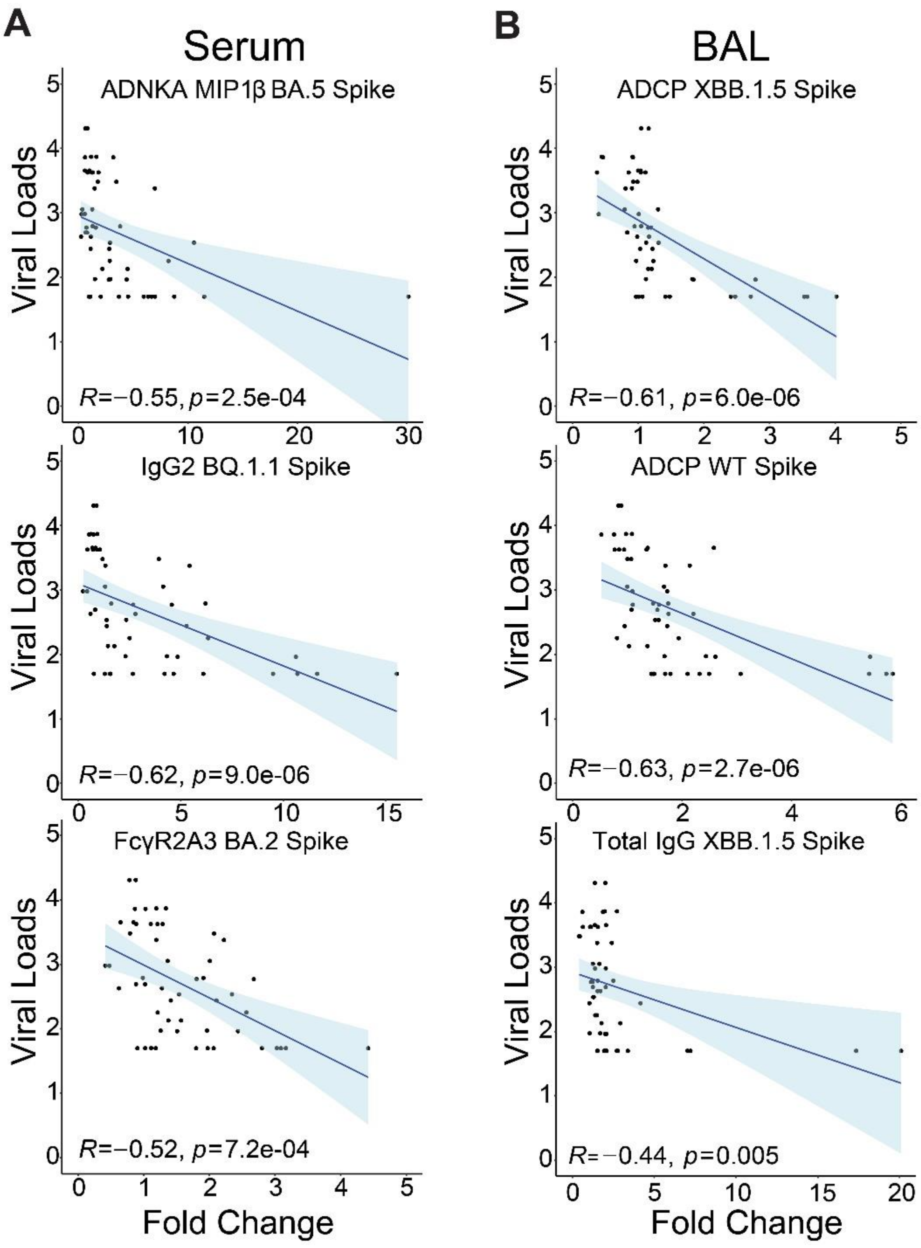


**Supplementary Figure 4. Validation of PLSR for selected correlates of protection by compartment**. A) Serum-based antibody correlates of protection selected by the PLSR model. Antibody feature measurements were plotted against viral loads and Spearman's R and p-values are shown for MIP1β, IgG2 to BQ.1.1 Spike, and FcγR2A3 to BA.2 Spike. On the x-axis is the fold change for the antibody feature against the viral loads, on the y-axis, for NHPs in all groups. B) Same as A, but for BAL-based antibody correlates of protection. Spearman's R and p-values are shown for ADCP to XBB.1.5 Spike, ADCP to WT Spike, and ADCD to BA.5 Spike. Spearman's R values and multiple comparisons adjusted p-values are shown.

**Supplementary Figure 5**

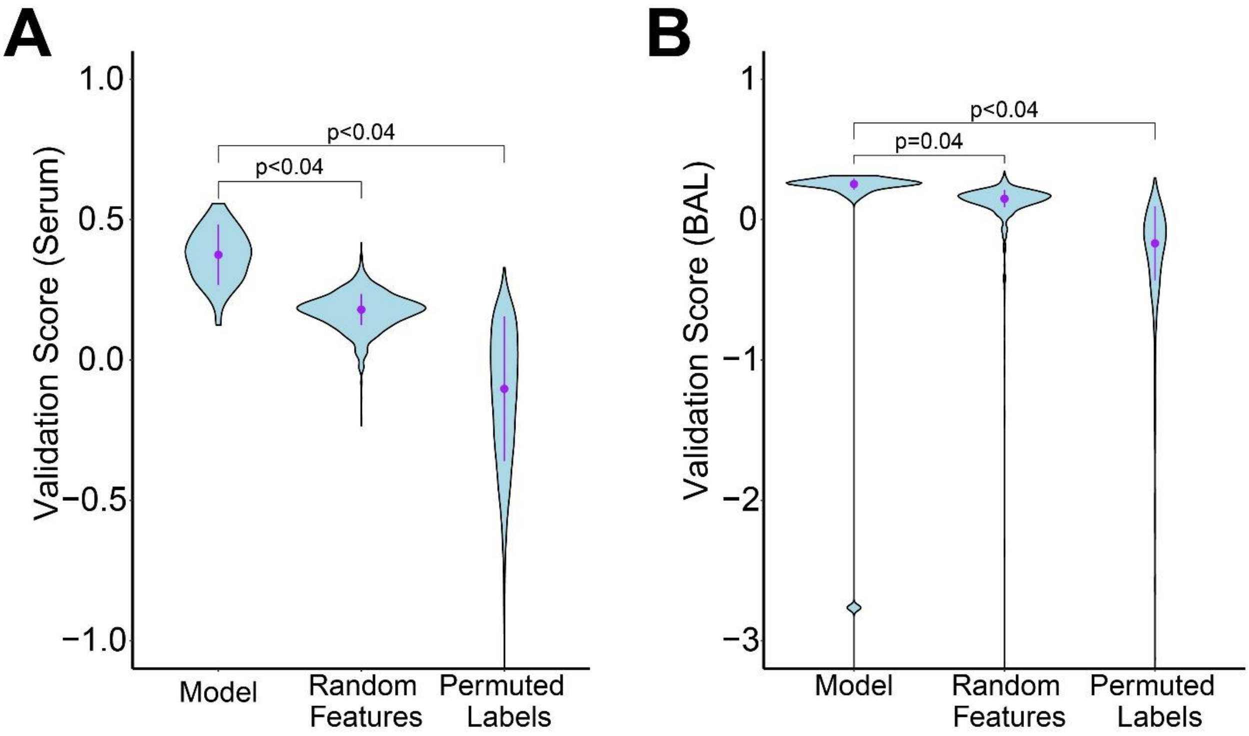


**Supplementary Figure 5. Validation of the PLSR models for Serum and BAL correlates of protection**. A) The PLSR model built to identify serum antibody correlates of protection was measured against random features and permutated labels. P-values for comparisons are shown above the violin plots. B) Same as A, but for the PLSR model built to identify BAL antibody correlates of protection.

**Supplementary Figure 6**

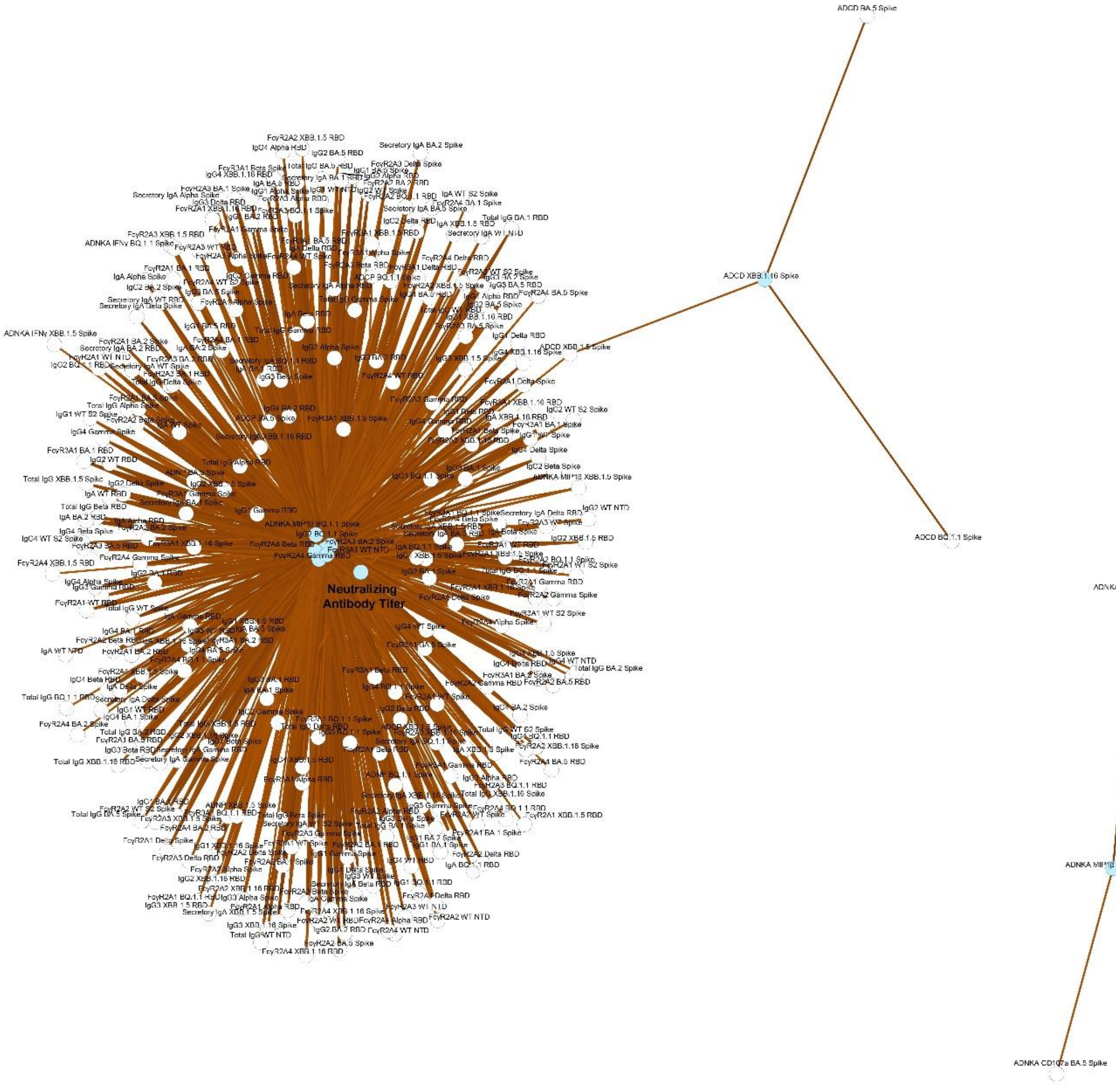


**Supplementary Figure 6. Full serum-based antibody correlates of protection constellation network**. Selected features driving protection in the serum selected by the PLSR and validated were plotted as nodes (blue circles) and significantly correlating features to each node are shown. Neutralizing antibody titer is shown in bolded letters in the constellation network. Positive correlations whose significance of correlation was < 0.05 after FDR correction are shown.

**Supplementary Figure 7**

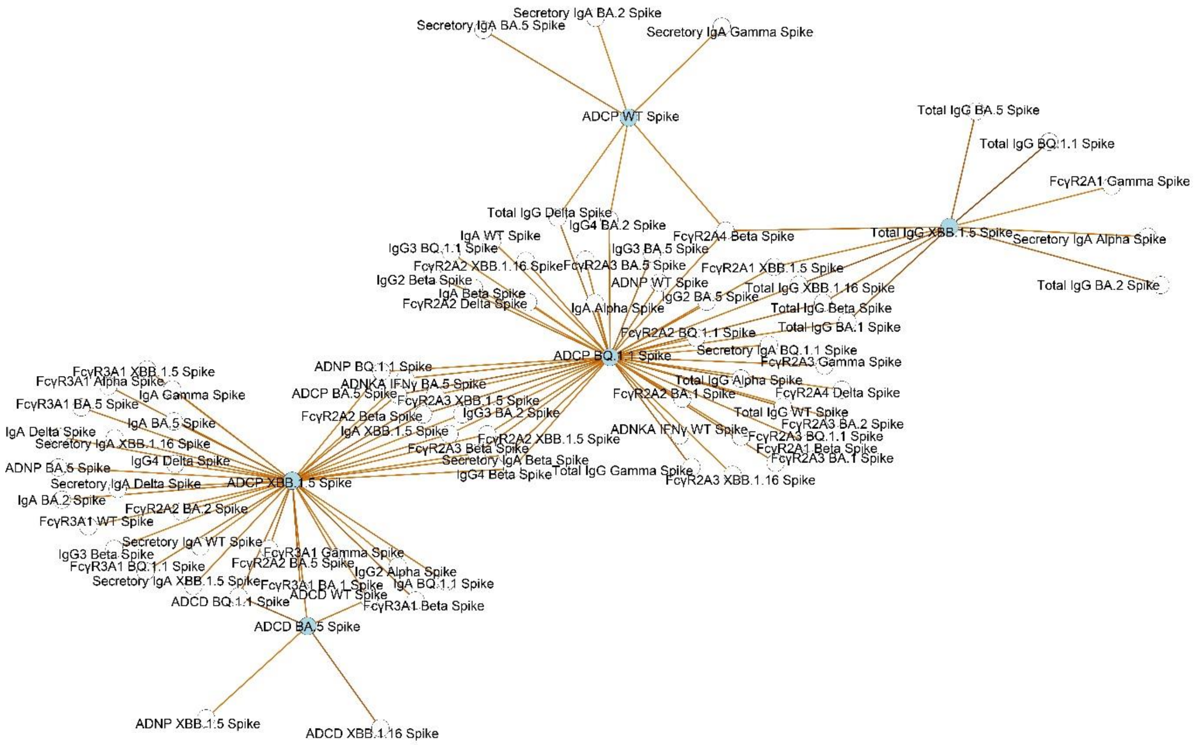


**Supplementary Figure 7. Full BAL-based antibody correlates of protection constellation network.** Selected features driving protection in the BAL selected by the PLSR and validated were plotted as nodes (blue circles) and significantly correlating features to each node are shown. Neutralizing antibody titer was not selected by the PLSR nor was it linked to the features. Positive correlations whose significance of correlation was $< 0.05$ after FDR correction are shown.

**Supplementary Figure 8**

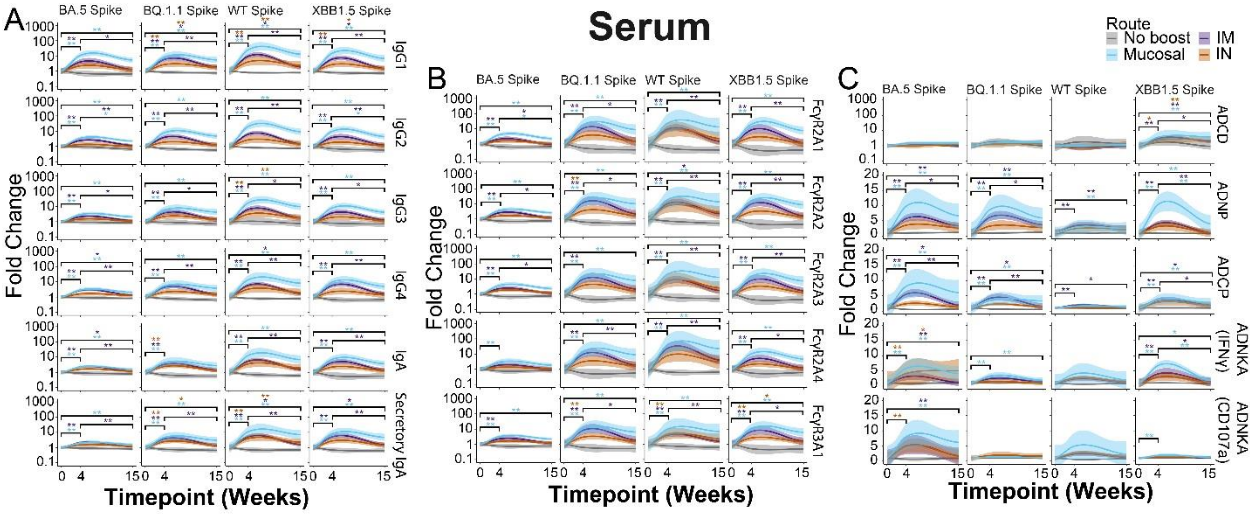


**Supplementary Figure 8. Univariate comparison of serum antibody-based binding and functions by booster delivery route**. A) Antibody isotype and subclasses binding were quantified against the indicated antigen by route of delivery. Shown are moving averages of serum antibody responses at week 0, week 4, and week 15 post-bivalent booster. Values are plotted as fold changes over week 0. B) Same as A, but for FcγR-binding antibodies. C) Antibody-based functional assays were quantified against the indicated antigen by route of delivery. Shown are ADCD, ADNP, ADCP, and ADNKA measured by INF-γ production and degranulation as measured by surface CD107a expression. Route of delivery color legend is shown in the top right corner. For all plots, * = $p < 0.05$, ** = $p < 0.01$; Wilcoxon Test followed by FDR correction for multiple comparisons.

**Supplementary Figure 9**

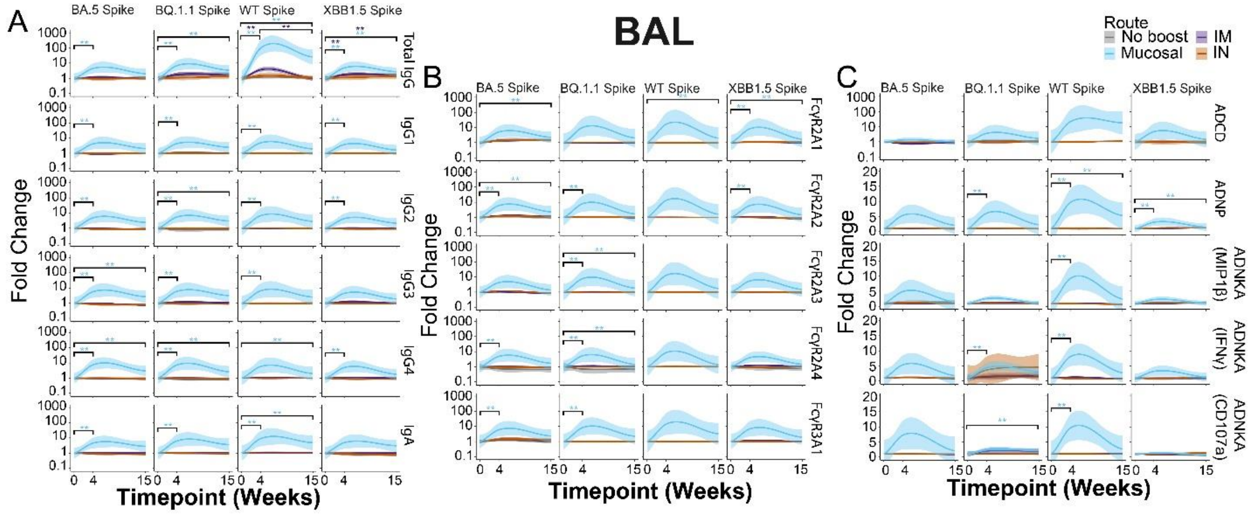


**Supplementary Figure 9. Univariate comparison of BAL antibody-based binding and functions by booster delivery route**. A) Antibody isotype and subclasses binding were quantified against the indicated antigen by route of delivery. Shown are moving averages of BAL antibody responses at week 0, week 4, and week 15 post-bivalent booster. Values are plotted as fold changes over week 0. B) Same as A, but for FcγR-binding antibodies. C) Antibody-based functional assays were quantified against the indicated antigen by route of delivery. Shown are ADCD, ADNP, ADCP, and ADNKA measured by INF-γ production and degranulation as measured by surface CD107a expression. Route of delivery color legend is shown in the top right corner. For all plots, * = $p < 0.05$, ** = $p < 0.01$; Wilcoxon Test followed by FDR correction for multiple comparisons.
