## Supplementary material for "Compartment-Specific Antibody Correlates of Protection to SARS-CoV-2 Omicron in Macaques": Key Resources Table

**Key Resource Table**

| **REAGENT or RESOURCE** | **SOURCE** | | **IDENTIFIER** |
| --- | --- | --- | --- |
| **Antibodies** |  |  |  |
| Anti-CD107a | BD Biosciences | 555802 | |
| Anti-CD3 | BD Biosciences | 558117 | |
| Anti-CD16 | BD Biosciences | 557758 | |
| Anti-CD56 | BD Biosciences | 557747 | |
| Anti MIP-1β | BD Biosciences | 550078 | |
| Anti-IFNγ | BD Biosciences | 340449 | |
| Anti-guinea pig complement C3 goat IgG fraction, FITC | MP Biomedicals | Cat # 0855385 | |
| Goat anti-Mouse IgG Fc Cross-Adsorbed Secondary Antibody, PE | Thermo Fischer | 31861 | |
| Mouse Anti-rhesus IgG-PE (SB108a) | Southern Biotech | 4700-09 | |
| Anti-rhesus IgG1 [7H11] | NHP Reagent Resource | 7H11 | |
| Anti-rhesus IgG2 [3C10] | NHP Reagent Resource | 3C10 | |
| Anti-rhesus IgG3 [2G11] | NHP Reagent Resource | 2G11 | |
| Anti-rhesus IgG4 [7A8] | NHP Reagent Resource | 7A8 | |
| Anti-rhesus IgA [9B9] | NHP Reagent Resource | 9B9 | |
| Anti-Secretory IgA | Invitrogen | MA1-81174 | |
| Anti-rhesus IgM | Life Diagnostics | 2C11-1-5 | |
| **Biological Samples** |  |  | |
| LowTox Guinea Pig Complement | CedarLane Labs | Cat # CL4051 | |
| **Chemicals, Peptides, and Recombinant Proteins** |  |  |  |
| SARS-CoV-2 WT Spike | Sino Biological | 40589-V08H4 | |
| SARS-CoV-2 BA1 RBD | Sino Biological | 40592-V08H129 | |
| SARS-CoV-2 WT S2 | Sino Biological | 40590-V08B | |
| SARS-CoV-2 WT NTD | Sino Biological | 40591-V49H | |
| SARS-CoV-2 Alpha Spike | Sino Biological | 40589-V08B6 | |
| SARS-CoV-2 Beta Spike | Sino Biological | 40589-V08B7 | |
| SARS-CoV-2 WT N | Sino Biological | 40588-V08B | |
| SARS-CoV-2 Delta Spike | Sino Biological | 40589-V08B16 | |
| SARS-CoV-2 Gamma Spike | Sino Biological | 40589-V08B10 | |
| SARS-CoV-2 XBB1.5 Spike | Sino Biological | 40589-V08H45 | |
| SARS-CoV-2 BA1 Spike | Sino Biological | 40589-V08H26 | |
| SARS-CoV-2 HCMV gB | Sino Biological | 10202-V08H1 | |
| SARS-CoV-2 XBB1.16 Spike | Sino Biological | 40589-V08H48 | |
| SARS-CoV-2 Alpha RBD | Sino Biological | 40592-V08H82 | |
| SARS-CoV-2 SARS Spike | Sino Biological | 40634-V08B | |
| SARS-CoV-2 Beta RBD | Sino Biological | 40592-V08H59 | |
| SARS-CoV-2 EG5 Spike | Sino Biological | 40589-V08H55 | |
| SARS-CoV-2 BA2.86 Spike | Sino Biological | 40589-V08H58 | |
| SARS-CoV-2 EG5 RBD | Sino Biological | 40592-V08H151 | |
| SARS-CoV-2 Delta RBD | Sino Biological | 40592-V08H91 | |
| SARS-CoV-2 Ebola Glycoprotein | Sino Biological | 40459-V08H | |
| SARS-CoV-2 Gamma RBD | Sino Biological | 40592-V08H86 | |
| SARS-CoV-2 BA2 Spike | Sino Biological | 40589-V08H28 | |
| SARS-CoV-2 BA2 RBD | Sino Biological | 40592-V08H123 | |
| SARS-CoV-2 BA5 Spike | Sino Biological | 40589-V08H32 | |
| SARS-CoV-2 BA5 RBD | Sino Biological | 40592-V08H131 | |
| SARS-CoV-2 BQ1.1 Spike | Sino Biological | 40589-V08H41 | |
| SARS-CoV-2 BQ1.1 RBD | Sino Biological | 40592-V08H143 | |
| SARS-CoV-2 XBB1.5 RBD | Sino Biological | 40592-V08H146 | |
| SARS-CoV-2 XBB1.16 RBD | Sino Biological | 40592-V08H136 | |
| Rhesus soluble Fcγ2A-1 | Duke University | Custom Order | |
| Rhesus soluble Fcγ2A-2 | Duke University | Custom Order | |
| Rhesus soluble Fcy2A-3 | Duke University | Custom Order | |
| Rhesus soluble Fcy2A-4 | Duke University | Custom Order | |
| Rhesus soluble Fcγ3A-1 | Duke University | Custom Order | |
| LC-LC-Sulfo-NHS Biotin | ThermoFisher | Cat # A35358 | |
| Brefeldin A | Sigma Aldrich | Cat # B7651 | |
| GolgiStop | BD Biosciences | Cat # 554724 | |
| Streptavidin-R-Phycoerythrin | Prozyme | Cat # PJ31S | |
| **Critical Commercial Assays** |  |  |  |
| Fix & Perm Cell Permeabilization Kit (Medium B) | ThermoFisher | GAS002S100 | |
| Fix & Perm Cell Permeabilization Kit (Medium A) | ThermoFisher | GAS001S100 | |
| EasySep™ Direct Human Neutrophil Isolation Kit | Stemcell Technologies | 19666 | |
| EasySep™ Human NK Cell Isolation Kit | Stemcell Technologies | 17955 | |
| NHS-Sulfo-LC-LC Kit | ThermoFisher | 21435 | |
| Zebra-Spin Desalting and Chromatography Columns | ThermoFisher | 89882 | |
| **Experimental Models: Cell Lines** |  |  |  |
| THP-1 monocytes | ATCC | RRID: CVCL_0006 | |
| Human Primary Natural Killer Cell Leukopack | StemCell | 200-0092 | |
| **Software and Algorithms** |  |  |  |
| GraphPad Prism 8 | GraphPad Software, Inc. | RRID:SCR_002798 | |
| iQue Forecyt 9.1 | Sartorius | 60028 | |
| R Studio V 6.0 | R Project for Statistical Computing | RRID:SCR_000432 | |
| Flow Jo | BD Bioscience | RRID:SCR_008520 | |
| MATLAB ver. R2019a | MathWorks | RRID:SCR_001622 | |
| **Other** |  |  |  |
| 384-well HydroSpeed Plate Washer | Tecan | 30190112 | |
| iQue Screener Plus | Intellicyt/Sartorius | 11811 | |
| MagPlex Microspheres | Luminex MFG | MC12001-01 (Cataloged by region) | |
| Green Fluorescent Neutravidin Microspheres | ThermoFisher | Custom Synthesis | |
| Red Fluorescent Neutravidin Microspheres | ThermoFisher | Custom Synthesis | |
| Scarlet Fluorescent Neutravidin Microspheres | ThermoFisher | Custom Synthesis | |
| 384-well HydroSpeed Plate Washer | Tecan | 30190112 | |
| MagPlex Microspheres | Luminex MFG | MC12001-01 (Cataloged by region) | |
| Green Fluorescent Neutravidin Microspheres | ThermoFisher | Custom Synthesis | |
| Red Fluorescent Neutravidin Microspheres | ThermoFisher | Custom Synthesis | |
| Scarlet Fluorescent Neutravidin Microspheres | ThermoFisher | Custom Synthesis | |
